# Behavioral prioritization enhances working memory precision and neural population gain

**DOI:** 10.1101/2021.09.16.460676

**Authors:** Aspen H. Yoo, Alfredo Bolaños, Grace E. Hallenbeck, Masih Rahmati, Thomas C. Sprague, Clayton E. Curtis

## Abstract

Humans allocate visual working memory (WM) resource according to behavioral relevance, resulting in more precise memories for more important items. Theoretically, items may be maintained by feature-tuned neural populations, where the relative gain of the populations encoding each item determines precision. To test this hypothesis, we compared the amplitudes of delay-period activity in the different parts of retinotopic maps representing each of several WM items, predicting amplitude would track with behavioral priority. Using fMRI, we scanned participants while they remembered the location of multiple items over a WM delay, then reported the location of one probed item using a memory-guided saccade. Importantly, items were not equally probable to be probed (0.6, 0.3, 0.1, 0.0), which was indicated with a pre-cue. We analyzed fMRI activity in ten visual field maps in occipital, parietal, and frontal cortex known to be important for visual WM. In early visual cortex, but not association cortex, the amplitude of BOLD activation within voxels corresponding to the retinotopic location of visual WM items increased with the priority of the item. Interestingly, these results were contrasted with a common finding that higher-level brain regions had greater delay-period activity, demonstrating a dissociation between the absolute amount of activity in a brain area, and the activity of different spatially-selective populations within it. These results suggest that the distribution of WM resources according to priority sculpts the relative gains of neural populations that encode items, offering a neural mechanism for how prioritization impacts memory precision.

## INTRODUCTION

Working memory (WM), the process involved in maintaining task-relevant information over a short period of time, links perception to behavior and is essential for a broad range of higher-level cognitive functions (Daneman & Carpenter, 1980; Engle et al., 1999; Süß et al., 2002). Multiple cortical areas are involved in maintaining information in working memory. In parietal and frontal areas, maintaining information in WM is characterized by sustained elevated neural activity during the delay period (Courtney et al., 1998; Curtis et al., 2004; Funahashi et al., 1989; Fuster & Alexander, 1971; McCarthy et al., 1994; Rowe et al., 2000). Furthermore, there is evidence that these brain areas are retinotopically organized (Jerde et al., 2012; Schluppeck et al., 2006), and voxels tuned to the location of a remembered stimulus show elevated delay-period activation compared to those tuned to other locations (Hallenbeck et al., 2021; Saber et al., 2015). Similarly, in retinotopic visual cortex, a location maintained in visual WM can be successfully decoded from delay period activity (Jerde et al., 2012; Rahmati et al., 2018; Sprague et al., 2014, 2016), along with other remembered visual features, like stimulus orientation, shape, motion direction, pattern, and/or color (Christophel et al., 2012; Harrison & Tong, 2009; Riggall & Postle, 2012; Serences et al., 2009, see Christophel et al., 2017 for a review). Interestingly, in these retinotopic regions, activation averaged across all voxels in a region does not typically display the sustained, elevated activation often seen in frontoparietal areas. (Christophel et al., 2012; Harrison & Tong, 2009; Riggall & Postle, 2012; Serences et al., 2009, but see Curtis & Sprague, 2021; Hallenbeck et al., 2021; Saber et al., 2015).

However, in these studies, participants maintained only a single item, or, multiple items of equal importance. In many cases, we need to remember multiple items with differing levels of importance, and thus, it would be beneficial to *prioritize* the representation of the most important information at the cost of a less robust representation of the least important information. Behavioral studies have demonstrated that participants can utilize cues about the relative reward and probe probability of different items in a WM display to flexibly prioritize the representation of the most valuable or likely-to-be-probed items (Bays, 2014; Emrich et al., 2017; Klyszejko et al., 2014; Yoo et al., 2018). In these studies, participants respond more accurately or precisely when asked to report the higher-priority items when compared to lower-priority items, suggesting that participants encode information in WM with varying levels of precision to accommodate task goals.

One plausible means of representing information in WM with varying levels of importance may be via modulating the strength of neural representations in a neural “priority map,” where the relative activation corresponding to different locations in the scene indexes the relative importance of that location (Bisley & Goldberg, 2010; Fecteau & Munoz, 2006; Jerde et al., 2012; Serences & Yantis, 2006; Thompson & Bichot, 2005). Indeed, there is extensive evidence that activation profiles across neural priority maps reflect the relative importance of different items in a display. For example, attention boosts the activity of neural populations with receptive fields that match the attended location, even in the absence of visual stimuli in the attended area (Buracas & Boynton, 2007; Gandhi et al., 1999; Gouws et al., 2014; Jerde et al., 2012; Kastner et al., 1999; Nobre et al., 2004; Rahmati et al., 2018; Saber et al., 2015; Serences & Yantis, 2007; Somers et al., 1999; Sprague et al., 2018)

Most of these studies examine the neural consequences of prioritized information on the selective attention of one item in a visual display, leaving unexplored how the brain prioritizes information held in WM (in the absence of visual input), and how multiple items are simultaneously represented across these neural priority maps. In this study, we test the hypothesis that activation in neural priority maps will have higher activation for items maintained in WM with higher priority, and this will in turn result in higher precision of WM reports. Furthermore, we hypothesize that any one neural priority map (one retinotopic area) can represent multiple items, and their relative priorities, through the activation of local populations tuned to each item’s location.

In our study, we asked if the response amplitude of neural populations during a working memory delay reflected the relative precision with which items were remembered. To answer this question, we collected event-related BOLD fMRI data from participants while they completed a multi-item spatial working memory task. In this task, each of the four items were pre-cued with a different probability of being probed for response, resulting in different memory precisions for each item (Yoo et al., 2018). Importantly, unlike previous research which only investigated the effects of attended versus unattended stimuli, this experimental design allowed us to simultaneously examine the effects of multiple levels of priority on neural amplitude. We used general linear models (GLMs) and population receptive field (pRF) mapping (Dumoulin & Wandell, 2008; Mackey et al., 2017) to estimate delay-period activity and location sensitivity, respectively, which allowed us to independently quantify location-specific delay-period activity for neural populations spatially tuned near the location of each item. We tested the prediction that higher priority items would exhibit higher delay-period BOLD activity at their corresponding retinotopic location in occipital (V1, V2, V3, V3AB), parietal (IPS0, IPS1, IPS2, IPS3), and frontal (iPCS, sPCS) brain areas. Remarkably, we find that this prediction holds in visual areas alone, suggesting prioritization of WM representations modulates neural gain. On the other hand, frontoparietal regions demonstrate clear elevated delay period activity, independent of the behavioral importance of different items. These results demonstrate a dissociation between the absolute activity in a brain area during a delay-period, and whether it encodes relative behavioral relevance through modulations of neural gain.

## EXPERIMENTAL METHODS

### Participants

Eleven participants (5 males, mean age=31.9, *SD*=6.8, 5 authors) participated in this experiment. Participants had normal or corrected-to-normal vision and no history of neurological disorders. Non-author participants were naive to the study hypotheses and were paid $30/hour. We obtained informed, written consent from all participants. The study was in accordance with the Declaration of Helsinki and was approved by the Institutional Review Board of New York University.

### Task procedures

We generated stimuli and interfaced with the MRI scanner, button-box, and eye-tracker using MATLAB software (The MathWorks, Natick, MA) and Psychophysics Toolbox 3 (Brainard, 1997; Pelli, 1997). Stimuli were presented using a PROPixx DLP LED projector (VPixx, Saint-Bruno, QC, Canada) located outside the scanner room and projected through a waveguide and onto a translucent screen located at the head of the scanner bore. Subjects viewed the screen at a total viewing distance of 64 cm through a mirror attached to the head coil. The display was a circular aperture with an approximately 32 degrees of visual angle (dva) diameter. A trigger pulse from the scanner synchronized the onsets of stimulus presentation and image acquisition.

Participants completed a multi-item probabilistic memory-guided saccade task (Fig. 1A). The fixation symbol in this experiment was an encircled fixation cross, with four equally-spaced concentric arcs within each quadrant. Each trial began with a 100 ms increase in the size of the outer circle of the fixation symbol. This was followed by a 700 ms endogenous precue which indicated the probe probability of each item. Probe probability was indicated through the number of illuminated arcs: all four arcs turned white in the quadrant corresponding to the 0.6 item, three arcs for the 0.3 item, two arcs for the 0.1 item, and zero arcs for the 0.0 stimulus. These probe probabilities were veridical across the entire experiment, though not necessarily for each block. The fixation pre-cue had a 0.6 dva radius around the center of the screen. The precue was followed by a 100 ms interstimulus interval, then by the items for 700 ms. The items were four white dots, one in each visual quadrant. Items were presented randomly between 9 and 10 dva from fixation. The location of the items in polar coordinates were pseudo-randomly sampled from every 10 degrees, avoiding cardinal axes. The item presentation was followed by a 10,100 ms delay. A response cue appeared afterward, which was a white wedge around the quadrant of the fixation symbol corresponding to the probed item. Participants made a memory-guided saccade to the remembered dot location within the corresponding quadrant of the screen. After the saccade, the actual dot location was presented as feedback and the participant made a corrective saccade to that location. After 800 ms, the feedback disappeared, participants returned their gaze to the central fixation cross, and a variable inter-trial interval began. The jittered inter-trial interval was pseudorandomly drawn from three time durations (8,800, 10,100 or 11,400 ms) to help deconvolve event-related activity associated with different trial epochs in the fMRI data. Each participant completed one scanning session consisting of 10-14 runs consisting of 12 trials each; they completed a total of 120-168 trials.

**Figure 1.**
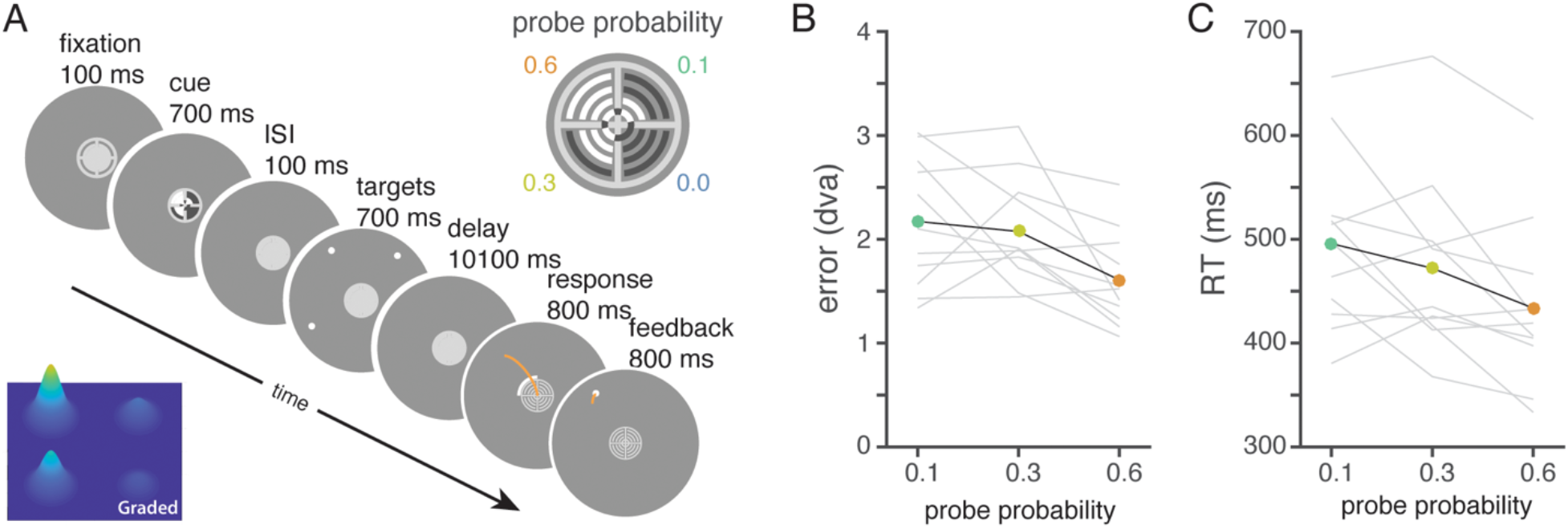
Prioritizing WM representations. *A*. Trial sequence. Participants viewed a precue that indicated the probe probabilities of the four targets, each presented in separate visual quadrants, by the number of arcs highlighted within the fixation symbol (top right inset). After the delay, one item was probed for response when a white arc appeared at the outer edge of one quadrant of the fixation symbol. Participants made a memory-guided saccade to the remembered location of the target. The true target location was then presented as feedback, which participants fixated. Bottom left inset: schematic predictions of priority map. Higher priority items will be represented with a taller bump. *B*. Memory error (grey: individual participants, black: mean) decreases with increasing priority (*b*=−1.16, *R^2^* = 0.18, *F*=6.67, *p*=.01) *C*. Memory-guided saccade response time (RT; grey: individual participants, black: mean) decreases with increasing priority, (*b*=−0.12, *R*^2^ =0.09, *F*=3.25, *p*=.08).

### Oculomotor methods

We recorded eye gaze data in the scanner at 1000 Hz (Eyelink 1000, SR Research, Ontario, Canada), beginning with a nine-point calibration and validation scheme. Using our freely available MATLAB iEye toolbox (github.com/clayspacelab/iEye_ts) we transformed raw gaze positions into degrees of visual angle, removed values outside of the screen area, removed artifacts due to blinks, smoothed gaze position with a Gaussian kernel with a standard deviation of 5 ms, and computed the velocity at each time point. Saccadic eye movements were defined with the following criteria: velocity ≥ 30 dva/s, duration ≥ 8 ms, and amplitude ≥ 0.25 dva. We define reaction time as the time between the response onset and the initialization of the first saccade, and error as the Euclidean distance between the target item and the last saccade landing position. For each trial, data were additionally drift corrected and calibrated to account for measurement noise, such that the gaze position during known trial epochs (i.e., fixation and response period) were at the correct location. Trials were excluded if the participant was not fixating during the delay period, no saccades were found during the response epoch, the initial saccade was too small in amplitude or too long in duration, or the final saccade error was greater than 10 degrees. These exclusion criteria resulted in removing between 4% and 51% (*M* = 23.9%, *SD* = 18.1%) of trials per participant. The total trial exclusions were especially high for four participants (42.5, 42.5, 47.5, 51.3%) because of poor eye-tracking quality, excessive sleepiness, and/or not making saccades during the response period. The exclusion rate of the remaining participants was *M* = 11.2%, *SD* = 5.0%. Removing the participants with high exclusion rates do not change the main effects in either the behavioral or neuroimaging results. We therefore retained the usable trials from these participants and performed all analyses with data from all participants.

### fMRI methods

#### MRI acquisition

All structural and functional MRI data were acquired on a 3T Siemens Prisma MRI system at the Center for Brain Imaging at New York University, using the CMRR MultiBand Accelerated EPI Pulse Sequences (Feinberg et al., 2010; Moeller et al., 2010; Xu et al., 2013). To acquire the functional BOLD contrast images, we used the following settings: Multiband (MB) 2D GE-EPI with MB factor of 4, 56 2-mm interleaved slices with no gap, voxel size 2mm, field-of-view (FoV) 208 × 208 mm, no in-plane acceleration, repetition time (TR) 1300 ms, echo time (TE) 42 ms, flip angle 66 deg, Bandwidth: 1924 Hz/pixel (0.64 ms echo spacing), posterior-anterior phase encoding, with fat saturation and “brain” shim mode. Distortion mapping scans, used to estimate the distortions present in the functional EPIs, were acquired with normal and reversed phase encoding after every other run. We used a 2D SE-EPI with readout matching that of the GE-EPI and same number of slices, no slice acceleration, TE/TR: 45.6/3537 ms, 3 volumes. We used T1-weighted MP-RAGE scans (0.8 × 0.8 × 0.8 mm voxels, 256 × 240 mm FoV, TE/TR 2.24/2400 ms, 192 slices, bandwidth 210 Hz/Pixel, turbo factor 240, flip angle 8 deg, inversion non-selective (TI: 1060 ms)) for gray matter segmentation, cortical flattening, registration, and visualization for creating ROIs.

### fMRI processing

#### Preprocessing

During preprocessing of functional data, we aligned the brain across runs and accounted for run-and session-specific distortions, with the aim of minimizing spatial transformations. This allowed us to maximize signal to noise ratio and minimize smoothing, ensuring data remains as near as possible to its original resolution. All preprocessing was done in Analysis of Functional NeuroImages (version: 17.3.09; Cox, 1996). First, we corrected functional images for intensity inhomogeneity induced by the high-density receive coil by dividing all images by a smoothed bias field (15 mm FWHM), computed as the ratio of signal in the receive field image acquired using the head coil to that acquired using the in-bore ‘body’ coil. Next, we estimated distortion and motion-correction parameters. To minimize the effect of movement on the distortion correction (the distortion field depends on the exact position of the head in the main field), we collected multiple distortion correction scans throughout the experiment. Thus, every two functional runs flanked the distortion scans used to estimate these parameters. We refer to the functional-distortion-functional scan as a mini-session. For each mini-session, we used the distortioncorrection procedure implemented in afni_proc.py to estimate parameters necessary to undistort and motion-correct functional images. Then, we used the estimated distortion field, motion correction transform for each volume, and functional-to-anatomical coregistration simultaneously to render functional data from native acquisition space into unwarped, motion corrected, and coregistered anatomical space for each participant at the same voxel size as data acquisition in a single transformation and resampling step. For each voxel on each run, we linearly detrended activation. We then computed percent signal change for each run.

### Estimating event-related BOLD activity

For each subject, we conducted a voxel-wise general linear model (GLM) to estimate each voxel’s response to different trial events: precue, stimulus, delay, and response. The BOLD activity of a single voxel was predicted from a convolution of a canonical model of the hemodynamic impulse response function and a box-car regressor which had length equal to each trial event.

Our main analysis investigates the effect of priority on the delay period. For all stimulus events that were not the delay period (precue, stimulus and response epochs), we used one predictor each to estimate their activity across trials. For the delay period, we defined a predictor for each trial, so that we could have single trial estimates of delay-period activity. The GLM additionally contained predictors to account for motion and intercept for each run, and were conducted in AFNI using 3dDeconvolve.

### Retinotopic mapping

We used a recently developed population receptive field (pRF) mapping approach (Mackey et al., 2017), which combines other pRF mapping approaches (Dumoulin & Wandell, 2008) with a more attentionally-demanding task in order to map topographic areas in occipital, parietal, and frontal cortex. The methods are briefly summarized below; a more detailed description can be found in (Mackey et al., 2017). During scanning, participants completed a difficult, covert attention task to ensure that required attention to the full spatial extent of the presented visual stimuli. The pRF mapping stimulus consisted of a bar subdivided into three random dot kinematograms (RDK; Williams & Sekuler, 1984; Fig. 2A). The participant indicated with a button press which of the two flanker rectangles contained dots moving in the same mean direction as the center rectangle. The bar (horizontal or vertical orientation) swept across the entire visual field throughout the experiment, so that participants had to attend to the areas of the visual field that contained visual stimuli. Each sweep lasted 31.2 s, and the bar updated its position every 2 TRs (2.6 s). Each run consisted of 12 sweeps, and participants completed between 9 and 12 identical runs, which were averaged together prior to model-fitting. Across sweeps, the width of the bar was varied among 3 discrete levels (1, 2, or 3 dva) to enable estimation of a nonlinear spatial summation model (Kay et al., 2013; Mackey et al., 2017).

**Figure 2.**
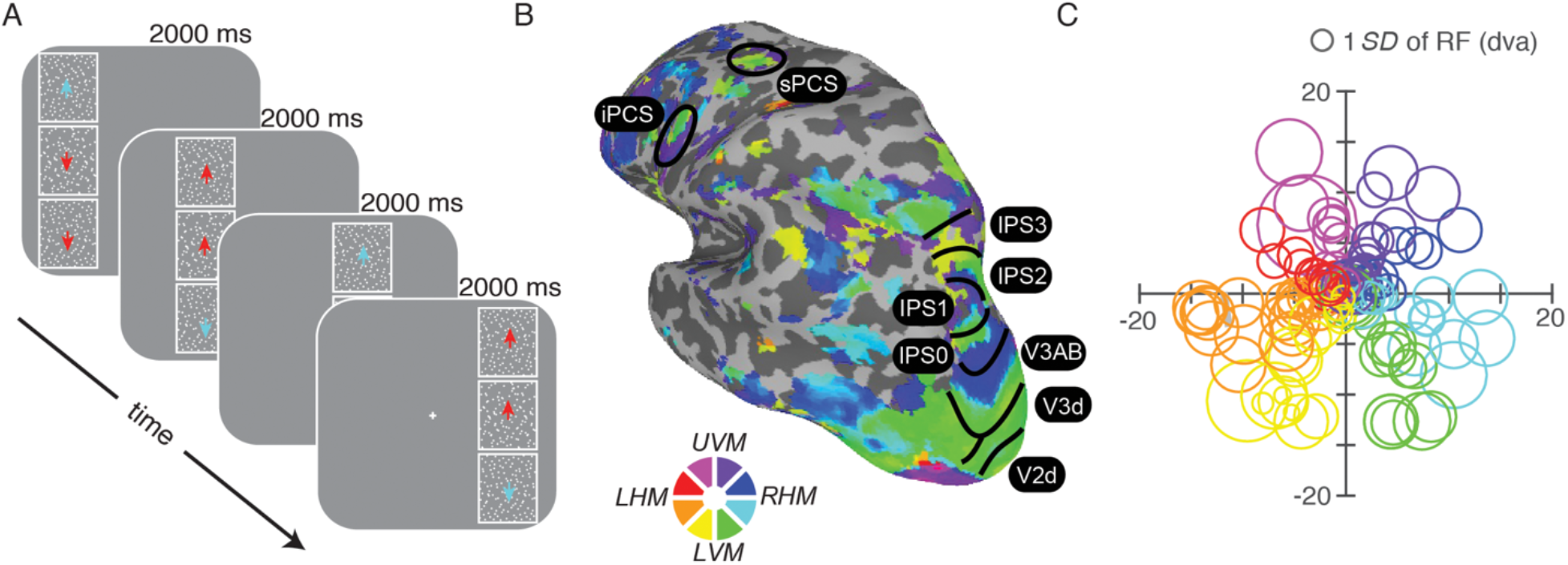
Measuring spatial selectivity of individual fMRI voxels via population receptive field mapping. *A*. Behavioral task used for pRF mapping. Participants indicated with a button press which of the two flanker rectangles contained dots moving in the same mean direction as the center rectangle. The rectangle configuration could sweep horizontally (as illustrated) or vertically. The dots within each rectangle moved orthogonal to the direction of the rectangle movement. Data from this task was used to fit pRF models to each voxel (nonlinear compressive spatial summation; Kay et al, 2013; Mackey et al, 2017). *B*. Example visual field maps plotted on an inflated surface view of the left cortex. The colors indicate the estimated polar angle center of each voxel’s pRF, which are used to define ROIs (UVM/LVM: upper/lower vertical meridian, LHM/RHM: left/right horizontal meridian). Colored voxels indicate pRF model fit explains ≥ 10% variance. *C*. Center position and standard deviation of 150 pRFs plotted from example participant’s V3. Like receptive fields of individual neurons, the receptive fields of voxels have less concentration and larger receptive fields with increasing retinal eccentricity.

We modeled the predicted response amplitude for each voxel at time *t*, 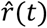using the following equation

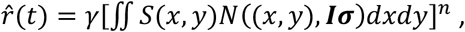

in which *S* is a binary stimulus image (1s where the stimulus was presented and 0s otherwise), and *N*((*x,y*),*Iσ*) is a Normal distribution with mean (*x, y*) and variance *Iσ*”, where *I* is a twodimensional identity matrix describing a circular, symmetric Gaussian. The parameters of this model are receptive field center (*x, y*), standard deviation σ, amplitude *γ*, and compressive spatial summation factor *n* (where *n* ≤ 1). Parameters were fit with a GPU-accelerated course grid search over parameters, followed by a local optimization method.

We used the estimated parameters acquired from the pRF mapping to define visual field maps. Specifically, we used the estimates of the polar angle and eccentricity of each voxel as measured through the pRF model. We visualized flattened cortical surface representations (computed using Freesurfer 6.0) of each subject’s brain using AFNI and SUMA (smoothed on the surface with a 5mm kernel), and defined retinotopic maps based on standard conventions (Larsson & Heeger, 2006; Mackey et al., 2017; Wandell et al., 2007). We defined the following areas: V1, V2, V3, V3AB, IPS0, IPS1, IP2, IPS3, iPCS, and sPCS (subset of LH regions visible in Figure 2B). In all further analyses, we use unsmoothed maps. First, we restricted our maps based on pRF estimates; we excluded voxels that did not have over 10% variance explained from the pRF model and voxels with RF centers nearer than 4 dva or farther than 20 dva eccentricity from fixation. We used a liberal exclusion criterion under the assumption that our analysis would essentially ignore voxels sensitive to locations far away from memorized item locations. Figure 2B illustrates the angle maps from an example participant, which demonstrates the topographic organization of visual field maps in occipital, parietal, and frontal cortical regions (Mackey et al., 2017).

### Analyses

We used two types of analyses to address our primary research questions. First, we analyzed whether average activation in each area exhibited sustained elevated delay period activity. Second, we tested whether the delay period activation associated with individual items differed based on their cued priority. While the first analysis asks whether the BOLD activity of an entire visual field map is elevated during the delay period, the second asks whether priority influences the *relative* profile of delay period activity within the map on each trial.

#### Univariate sustained activation analysis

First, we tested whether each visual field map exhibited elevated delay-period activity. For each participant in each visual field map, we averaged all the delay-period predictors, which we denote β, across trials and voxels to get one estimate of delay-period activity. For each ROI, we evaluated statistical significance using parametric t-tests (average β across participants compared against 0, one-tailed) as well as nonparametrically through bootstrapped confidence intervals (resampling the 11 participants’ average betas, with replacement, a thousand times and comparing the proportion of resampled means below 0).

#### Item-specific delay-period activation

Second, we tested whether the delay-period activity varied based on cued priority associated with each individual item. The goal of this analysis is to test whether the delay-period activity was different for populations maintaining items at different priority levels. While the previous analysis investigates if an entire visual field map has sustained activity through a delay period, this analysis tests whether the relative activity within a visual field map differs across different levels of priority.

To dissociate the delay-period activity evoked by different items, we estimated item-specific population activities by adjusting the contribution of each voxel’s delay-period activity (estimated through GLM) according to its selectivity for each item’s location. This means that the larger the overlap between a voxel’s pRF and an item’s location, the larger the contribution of that voxel in the overall activity evoked by that item. For every item in every trial, we computed a pRF-weighted β,

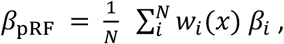

where *w_i_*(*x*) is the weight associated with the β^th^ voxel at location *x*, and β_i_ is the GLM-acquired delay-period β at voxel *i*. We define *w_i_* as the receptive field of voxel _i_, which we model in accordance with the pRF models; voxel *i*. ‘s receptive field is represented as a non-normalized Gaussian with mean *μ_i_* and variance σ_i_^2^,

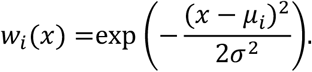

Thus, voxels have a higher weight when they are “more tuned” to an item’s location. For example, when an item is at the voxel’s receptive field center, or when *x* = *μ_i_*, the weight is *w_i_*(*x*) = 1. As the Euclidean distance between *x* and *μ_i_* increases, this weight decreases; the steepness with which it decreases is related to σ_i_. This weighting is mathematically similar to pRF-weighted “stimulus reconstructions” reported previously (Kok & de Lange, 2014; Thirion et al., 2006), but evaluated at single points of the image (the item positions).

For each visual field map, we test a significant effect of priority. First, for each participant, we conducted a linear regression with *β*_pRF_ as the dependent variable, priority level as the predictor. Then, we conducted a t-test across participants to see if the priority predictor was above 0.

## RESULTS

### Behavioral results

To test our prediction that higher priority items would be remembered more precisely, we looked at memory error as a function of priority. We operationalized error as the Euclidean distance between the target location and the final saccade landing point. We conducted a linear regression testing whether error varied based on priority. We found that increasing priority significantly predicted a decrease in error (*b*=−1.16, *R^2^* = 0.18, *F*=6.67, *p*=.01). This result was not due to a speed-accuracy trade off, because reaction times also marginally decreased with increasing priority (*b*=−0.12, *R*^2^=0.09, *F*=3.25, *p*=.08). These results indicate that people remember higher priority items more precisely, consistent with previous reports (Bays, 2014; Emrich et al., 2017; Klyszejko et al., 2014; Yoo et al., 2018).

### Neuroimaging results

We investigated the effect of priority and working memory over 10 retinotopically-defined brain regions across dorsal visual, parietal, and frontal cortex: V1, V2, V3, V3AB, IPS0, IPS1, IPS2, IPS3, iPCS, and sPCS. For each brain area, we asked whether there was elevated delay activity in the entire region, then whether there was an item-specific effect of priority on the magnitude of delay period activity within the region. For all statistics reported, *p*-values were Bonferroni corrected across ROIs.

### Univariate delay-period activation

First, we tested whether any retinotopic regions showed elevated delay period activity when activation was averaged across voxels within the region. Delay-period activity was quantified using a single-trial GLM approach (see Methods). Early visual areas V1, V2, and V3 did not exhibit activity that was significantly higher than baseline (Figure 4A, 6). However, dorsal visual region V3AB and all parietal and prefrontal regions tested did exhibit delay-period activity that was significantly higher than baseline (Fig. 5A, 6; *p*<.001, for t-test and bootstrapped significance test).

### Item-specific delay-period activation

Second, we tested whether the delay-period BOLD activity in voxels spatially selective for locations of different items differed based on the item’s behavioral priority. We used each voxel’s independently-estimated population receptive field (pRF) to quantify the overall population activation associated with each item’s spatial location on each trial (Fig. 3; see Methods). This results in a measured activation level corresponding to each WM item (0.6, 0.3, 0.1, 0.0 probe probability) on each trial (Fig. 4B-C; 5B-C). We consider this analysis a higher-resolution investigation of the effect of the behavioral priority of multiple items on the measured priority map within an ROI. To establish whether the cued priority significantly sculpted activation strength in each ROIs retinotopic map, we first conducted a linear regression between the behavioral priority level and the item-specific delay period activity computed based on *β*_pRF_ for each participant (see Methods). We then conducted a t-test to investigate if the priority significantly predicted *β*_pRF_ with slope above 0.

**Figure 3:**
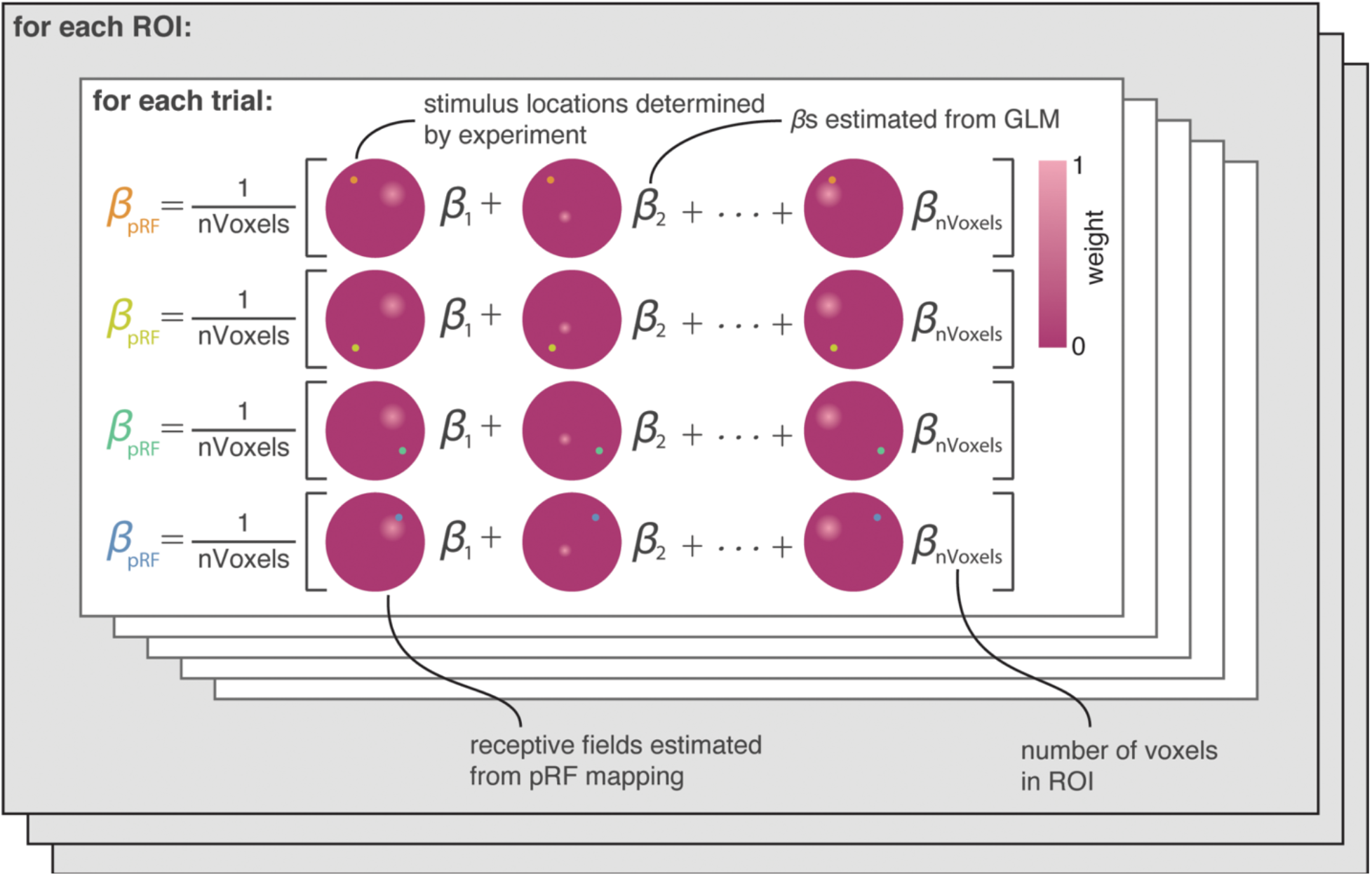
Procedure for estimating WM representation gain (*β*_pRF_) based on pRF parameters. For each trial, we weight the GLM-obtained estimate of delay-period activity for voxel *i*, *β_i_*, by its sensitivity to the location of the current item as estimated by the pRF model. The heat plots illustrate the estimated receptive field over the entire stimulus display (a non-normalized Gaussian distribution with a mode equivalent to weight = 1) for each voxel. The corresponding weight for each voxel is the scalar value at the item’s location (the value at the colored dot). Each row here illustrates how the weights are updated across items within the same trial by changing the item’s location (the colored dots on the receptive field maps). Across trials, the βs and stimulus locations are updated. Across ROIs, the voxels and their receptive fields are updated.

**Figure 4.**
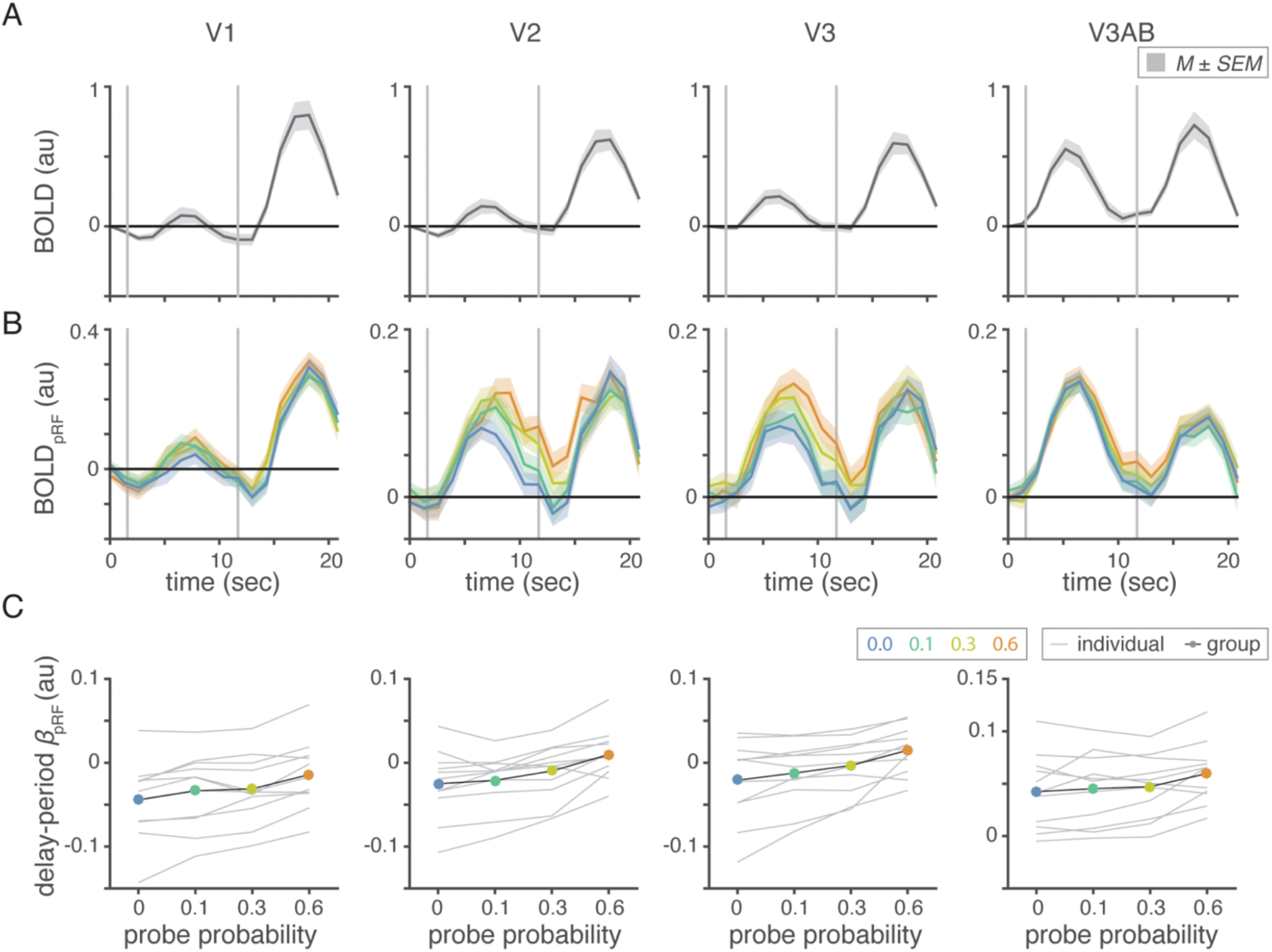
Delay-period activity in visual cortex increases with priority. Univariate HRF timecourses (A), *β*_pRF_-weighted HRF timecourses (B), and item-specific delay-period activation (C) for visual areas V1, V2, V3, and V3AB (columns). *A*. Average trial-related BOLD signal (*M* ± *SEM* across participants), independent of priority showing event-related changes in BOLD activity. Univariate delay-period activation does not have elevated delay-period activity. *B*. Average pRF-weighted BOLD signal (*M* ± *SEM* across participants). Apparent separation of BOLD activity during timecourse corresponding to HRF of delay period. *C*. *β*_pRF_as a function of priority for individual participants (grey) and across participants (black). *β*_pRF_increases with priority.

**Figure 5.**
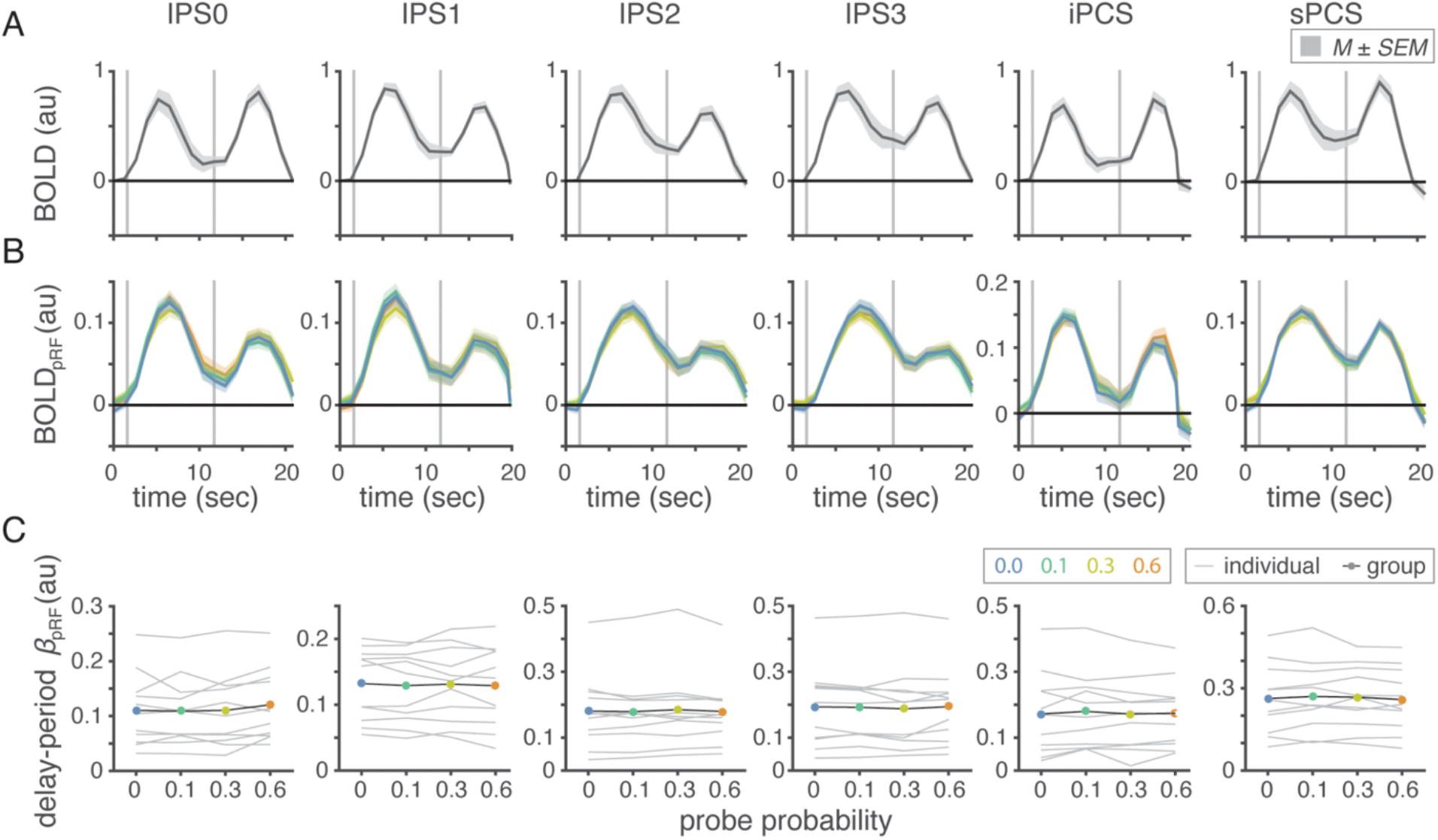
No effect of priority in delay activity in parietal and frontal areas. Univariate HRF timecourses (A), *β*_pRF_-weighted HRF timecourses (B), and item-specific delay-period activation (C) for parietal and frontal areas IPS0, IPS1, IPS2, IPS3, iPCS, and sPCS (columns). *A*. Average trial-related BOLD signal (*M* ± *SEM* across participants), independent of priority showing event-related changes in BOLD activity. Delay period shows an elevated delay-period activity. *B*. Average pRF-weighted BOLD signal (*M* ± *SEM* across participants). No apparent separation of BOLD activity during timecourse corresponding to HRF of delay period. *C*. *β*_pRF_as a function of priority for individual participants (grey) and across participants (black). There is no effect of priority on *β*_pRF_.

While there was no overall elevated delay-period activation in visual areas (Fig. 4A; 6A), the profile of activation within each region showed an effect of priority (Fig. 4B-C; 6B). Priority was a significant modulator of delay-period BOLD activity for V1 (*t*(10)=6.47, *p*=.0007), V2 (*t*(10)=4.74, *p*=.007), and V3 (*t*(10)=4.13, *p*=.02). Area V3AB was significant before Bonferroni correction, but trending afterward (*t*(10)=3.45, *p*=.06). These results suggest that subpopulations in visual areas corresponding to an item’s location have higher delay-period activity when maintaining a higher priority item (Figure 4B). On the other hand, there was no effect of priority on item-specific delayperiod activity in any frontoparietal region (Figure 5B-C; 6B).

**Figure 6.**
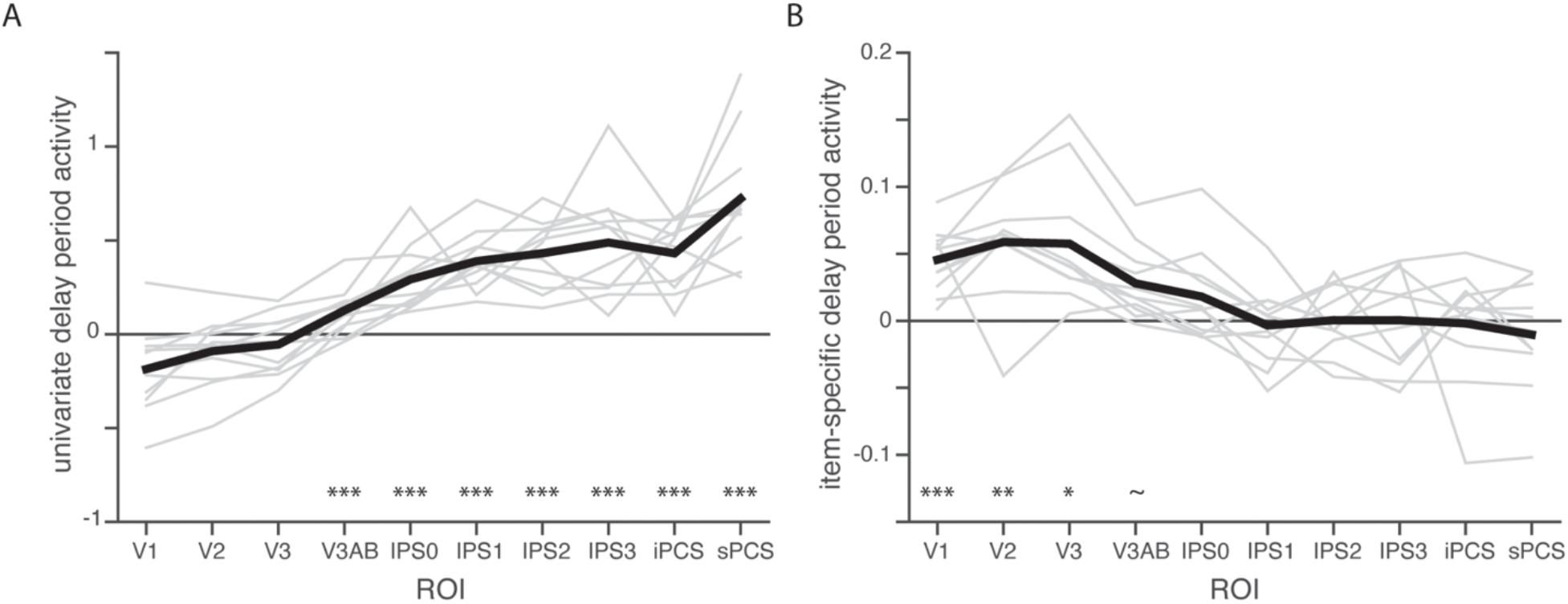
Summary of fMRI results. *A*. Univariate delay-period activity for each ROI, for individual participants (grey lines) and averaged across participants (black line) as estimated from GLM. Asterisks indicate ROIs that have delay-period activity significantly higher than baseline after Bonferroni correction (*** p<.001). *B*. Estimated slope of pRF-weighted delay-period activity for individual participants (grey lines) and averaged across participants (black line). Asterisks indicate ROIs that have a significant effect of behavioral priority on item-specific delay-period activity after Bonferroni correction. (*** p<.001, ** p<.01, * p<.05, ~ p<.10).

## DISCUSSION

In this study, we asked how the behavioral relevance of an item in working memory modulates neural representations across the brain. We hypothesized that an item’s priority would be reflected in the gain of the same neural subpopulations that encode its location. To test this hypothesis, we collected fMRI BOLD activity while participants completed a multi-item spatial delayed estimation task and analyzed the delay period activity using traditional univariate metrics and a novel method informed by computational modeling of voxel receptive field properties. We found two main results: first, frontoparietal areas exhibited elevated delay-period BOLD activity on average, but did not exhibit item-specific gain as a function of priority; second, visual areas exhibited item-specific gain as a function of priority but no elevated delay-period BOLD activity on average. The dissociation between univariate activity and item-specific activity suggests different roles across the processing hierarchy. What are these distinct roles, and how do they relate to WM theory?

Our results in visual areas are consistent with decoding studies finding successful decoding of stimulus-information from visual cortex despite a lack of sustained delay-period activity (Albers et al., 2013; Christophel et al., 2012; Ester et al., 2009, 2015; Harrison & Tong, 2009; Lee et al., 2013; Riggall & Postle, 2012; Serences et al., 2009; van Bergen et al., 2015). While we did not aim to decode specific item positions, our observation that activation in retinotopic ROIs in voxels tuned near high-priority items exhibit greater delay-period activation than voxels tuned to lower-priority items is consistent with a mechanism whereby a prioritized item in WM is maintained with higher gain. Because our study made the very specific hypothesis that precision depends on the gain of the same neural subpopulations encoding an item’s location during the WM delay period, our results offer an explanation beyond previous decoding studies by explaining *how* behavioral priority impacts neural codes, not merely *if* it has an effect.

A promising explanation for the role of sensory areas in WM comes from the sensory recruitment hypothesis, which posits that the same areas that process sensory information are also involved in the maintenance of sensory WM representation (Curtis & D’Esposito, 2003; Curtis & Sprague, 2021; Pasternak & Greenlee, 2005; Postle, 2006; Serences, 2016). One appealing aspect of this hypothesis relates to efficiency in that WM representations are precisely maintained in the same regions using the same mechanisms used for encoding percepts. Our results also appeal to efficiency as they demonstrate that the early visual areas that have been repeatedly shown to encode WM features also encode the relative priority of multiple WM items.

Previous studies have tested how changing an item’s priority status over the course of a trial impacts the quality of decoded neural representations from visual, parietal, and frontal cortex (Christophel et al., 2018; Iamshchinina et al., 2021; LaRocque et al., 2017; Lewis-Peacock et al., 2012; Lorenc et al., 2020; Rahmati et al., 2018; Rose et al., 2016; Sahan et al., 2020; Sprague et al., 2016; Yu et al., 2020). In many of these studies, experimenters have employed a ‘dual retro-cue’ task, in which cues presented during the delay period allow participants to transiently prioritize one of multiple items for an upcoming response. Importantly, when an item is cued, the cue is often 100% valid, and so any non-cued items can be transiently deprioritized. Thus, these tasks do not test how multiple items of continuously-varying priority states are represented simultaneously. However, they have the unique ability to compare coding properties of WM representations imminently relevant for behavior to those relevant at a later timepoint. Results from these studies vary and suggest that only immediately-relevant WM representations can be decoded (Lorenc et al., 2020; Sahan et al., 2020; Yu et al., 2020), that both prioritized and non-prioritized WM representations can be decoded with sufficient sample size (Christophel et al., 2018), or that these results critically depend on particulars of the multivariate analysis procedures employed (Iamshchinina et al., 2021). Using an antisaccade procedure, where the goal of the memory-guided saccade was opposite the visual target, we recently demonstrated that the strength of WM representations in early visual cortex migrated from the visual target early in the delay to the saccade goal later in the delay (Rahmati et al., 2018). In light of the current results, we argue that the priority of the two locations changed during the course of the trial as the location of the visual stimulus was recoded into the location of the memory-guided saccade. Indeed, future studies employing dynamic reprioritization of spatial positions throughout a trial (e.g., Sprague et al., 2016) may help triangulate how graded relative priority and transient reliable priority compare to one another.

The relationship between behavioral priority and neural gain additionally provides support for probabilistic population coding (PPC) models, which predict that increased neural gain increases SNR in neural populations, resulting in higher behavioral precision (Bays, 2014). PPC models posit that neural populations themselves represent a probability distribution over a stimulus representation, such that increased neural response gain corresponds to more precise representations of that stimulus (Ma et al., 2006; Zemel et al., 1998). In working memory tasks, population coding models have used neural gain as a proxy for the memory precision of items, and are able to account for the effects of prioritization on memory error (Bays, 2014; Ma et al., 2006; Seung & Sompolinsky, 1993). Our experimental design extends results of previous studies by simultaneously recording items of multiple levels of priority on every trial, and showing that the delay-period activity increases with cued priority in a graded fashion, which tracks with changes in behavioral memory errors (Fig. 1B). These results are also consistent with results finding decoded error tracks memory error (Li et al., 2021) and decoded uncertainty decreased with increasing behavioral priority (Levin et al., 2021). Perhaps the sensory recruitment hypothesis and PPC together provide an explanation for the cortical specificity of our results; only these posterior areas code sensory information with high enough resolution to see differences in item-specific gain.

Our results in frontoparietal areas are especially interesting in reference to decoding studies which typically find an absence of stimulus information in frontoparietal regions despite elevated delay-period activity (e.g., Christophel et al., 2012; Emrich et al., 2013; Postle et al., 2003; Riggall & Postle, 2012, but see Ester et al., 2015; Hallenbeck et al., 2021; Jerde et al., 2012 for conflicting results). Because our study made the very specific hypothesis that precision is represented through the gain of the same neural subpopulations encoding an item’s location during the WM delay period, unlike decoding studies, we cannot conclude that priority is not represented in frontoparietal areas at all.

These findings, however, are consistent with a larger body of literature demonstrating that frontal areas have been considered to play a more goal-oriented role. For example, Riggall and Postle (2012) were able to decode task relevant information (i.e., stimulus dimension) but not stimulus information (i.e, specific stimulus feature) in frontal cortex (but see Ester et al, 2015). Perhaps PFC encodes task demands more than low-level physical properties of stimuli (Christophel et al., 2012, 2015; Christophel & Haynes, 2014; Emrich et al., 2013; Freedman et al., 2002; Freedman et al., 2001; McKee et al., 2014; Riggall & Postle, 2012; Wallis et al., 2001; Wallis & Miller, 2003; Warden & Miller, 2010; White & Wise, 1999).

In summary, our study adds to the growing body of literature connecting activation patterns in visual areas to high-fidelity WM representations in the brain. By applying a novel pRF-guided analysis method, we demonstrated that voxels tuned to higher-priority locations held in WM exhibited greater gain during the delay period, consistent with a higher-precision representation, which we observed behaviorally. Modulating neural gain may be a key mechanism whereby neural systems dynamically optimize processing resources to support cognitive demands.

## Acknowledgements

We thank the Center for Brain Imaging at NYU and their staff for support during data collection. This research was supported by the National Eye Institute (R01 EY-016407 and R01 EY-027925 to C.E. Curtis, F32 EY-028438 to T.C. Sprague, and T32EY007136-27 to New York University & G.E. Hallenbeck), a Sloan Research Fellowship to T. C. Sprague, and an NVidia Hardware Grant to T.C. Sprague and C.E. Curtis.

